# Stable isotope tracer captures the anabolic response of human skeletal muscle microtissues undetected by puromycin labeling

**DOI:** 10.1101/2025.09.12.675841

**Authors:** Yekaterina Tiper, Cassidy T Tinline-Goodfellow, Penney M Gilbert, Daniel R Moore

## Abstract

Skeletal muscle microtissues are valuable *in vitro* models for studying the stimuli regulating muscle protein synthesis (MPS), the key determinant of changes in muscle mass. Differentiated between opposing posts, microtissues contain aligned, contractile myotubes, providing a controlled system for investigating the responses of skeletal muscle to nutrient and contractile stimulation. However, microtissue MPS responses to these stimuli remain under-characterized. Stable isotope-labeled amino acid tracers deliver sarcoplasmic and myofibrillar fractional synthetic rates (FSR) for MPS in human studies, but have not been implemented in engineered skeletal muscle. We close these gaps by characterizing stimulation-induced MPS, in microtissues and 2D myotubes derived from the same primary myoblast line, using stable isotope tracers and puromycin incorporation. In microtissues, sarcoplasmic FSR increased significantly during the two-hour period following amino acid treatment (*p* < 0.0001), whereas myofibrillar FSR remained unchanged (*p* = 0.159). However, both fractions were unresponsive to ketone stimulation and contraction (all *p* ≥ 0.703). 2D myotubes showed significant increases in sarcoplasmic and myofibrillar FSR in response to amino acid treatment (both *p* = 0.002). Notably, microtissues demonstrated a more stable myofibrillar protein fraction, with a sarcoplasmic-to-myofibrillar FSR ratio of ∼2:1 which closely resembled that of native human muscle. The puromycin-based approach failed to detect MPS responses to any stimulus (all p ≥ 0.677), highlighting the superior sensitivity of tracer-based measurements, particularly where longer timescales are needed to capture an effect. These findings support the use of engineered muscle and isotope-derived measurements of MPS in future studies of stimuli regulating skeletal muscle mass.

**New Findings:** *What is the central question of this study?:* Stable isotope tracers are emerging as a powerful approach to measure fraction-specific protein synthesis. However, their efficacy relative to conventional puromycin labeling remains unreported, and they have not been applied to engineered skeletal muscle.

*What is the main finding and its importance?:* By implementing stable isotope tracers in engineered muscle, we showcase the ability to capture anabolic responses that are undetected by puromycin-based methods. We found that the myofibrillar protein fraction of microtissues is more stable than the sarcoplasmic fraction, a property of native muscle, absent in 2D myotubes. These findings demonstrate the physiological relevance of engineered muscle and support the adoption of isotope-derived measurements in future studies.

## 1. INTRODUCTION

Skeletal muscle plays a critical role in overall health, facilitating movement (Mukund & Subramaniam, 2020), whole-body protein metabolism (Wolfe, 2006), and glucose homeostasis (Merz & Thurmond, 2020). Maintaining muscle mass throughout life is essential, as age-related muscle loss, or sarcopenia, can lead to increased frailty and a higher risk of falls and fractures (Yeung et al., 2019). Muscle protein synthesis (MPS) is the primary determinant of net muscle balance, ultimately determining muscle net balance, more so than muscle protein break down. The most potent anabolic stimuli of MPS are protein ingestion and resistance exercise (Moore, 2019). In recent years, ketone bodies, particularly β-hydroxybutyrate and acetoacetate, have also gained increasing interest for their role in enhancing MPS. It has been reported that ketone monoester ingestion in the overnight post-absorbative state potentiates MPS to a level comparable to that of a moderate protein intake (Hannaian et al., 2024).

Studies using two-dimensional (2D) myotube cultures *in vitro* have advanced our understanding of the molecular mechanisms that regulate skeletal muscle mass, particularly the balance between MPS and MPB (Gran & Cameron-Smith, 2011; Huang et al., 2007; Rommel et al., 2001). These models eliminate many of the confounding factors present *in vivo*, such as fluctuating hormone levels and immune system responses, while offering greater experimental control over variables like dosage, nutrient availability, and treatment duration. Additionally, they enable higher-throughput experimentation due to their simpler handling and scalability compared to *in vivo* studies. However, 2D myotube cultures are limited in their ability to model native muscle structure and function, as they lack alignment and tend to detach from the substrate upon contraction. To overcome these limitations, three-dimensional (3D) models of skeletal muscle have emerged (Hofemeier et al., 2021; Madden et al., 2015; Vandenburgh et al., 2008). These models involve the differentiation of myoblasts in an extracellular matrix under uniaxial tension between two fixed points. Unlike 2D systems, 3D skeletal muscle models reproduce key aspects of native muscle architecture, support prolonged culture durations, and facilitate studies of contractile stimuli, while retaining the advantages of *in vitro* experimentation (Khodabukus, 2021). To fully harness the potential of 3D models in investigating skeletal muscle mass regulation, reliable methods for measuring MPS in these models are required.

Puromycin is a widely used tool to measure MPS rates *in vitro*. This aminonucleoside antibiotic, produced by the bacterium *Streptomyces alboniger*, acts as a structural analog of aminoacyl tRNAs. It incorporates into nascent peptide chains but prematurely terminates their elongation due to its non-hydrolyzable amide bond. When used in minimal amounts over 5-30 min pulses, puromycin incorporation into newly synthesized proteins reflects the rate of mRNA translation, which can be semi-quantitatively assessed via immunoblotting of cell lysates (Schmidt et al., 2009). This technique, known as surface sensing of translation (SUnSET), is widely used in skeletal muscle research to measure protein synthesis both *in vitro* and in animal models (Goodman et al., 2011).

Due to the toxicity of puromycin, it is unsuitable for human use. *In vivo* human studies of MPS commonly rely on stable isotope-labeled amino acid tracers (e.g., L-*[ring-*^*2*^*H*_*5*_]-phenylalanine, L-[1-^13^C]-leucine, etc.) administered orally or intravenously and quantitatively analyzed via gas or liquid chromatography-mass spectrometry (Wilkinson et al., 2017). Stable isotope tracers, being chemically and functionally identical to their naturally occurring counterparts, allow amino acid kinetics to be traced over extended periods through the behaviour of the labeled compounds. In the tracer incorporation model, the rate of tracer incorporation into newly synthesized peptides is quantified over time, yielding fractional synthetic rates (FSR) as a measure of MPS (Kim et al., 2016). FSR can be assessed as a mixed protein fraction (Moore, Robinson, et al., 2009) or as myofibrillar and sarcoplasmic protein fractions (Moore, Tang, et al., 2009). Studies investigating fraction-specific MPS have demonstrated that the contractile myofibrillar pool is sensitive to both resistance exercise and amino acid ingestion, while the sarcoplasmic pool primarily responds to amino acid ingestion (Moore, Tang, et al., 2009).

Stable isotope-labeled amino acid tracers have been applied to 2D myotube systems (e.g., Crossland et al., 2020) but have yet to be utilized in engineered skeletal muscle. Consequently, fraction-specific MPS has not been investigated in 3D systems over extended periods under conditions of nutrient availability and contractile stimulation. We previously developed MyoTACTIC, a 96-well culture platform for the fabrication of human skeletal muscle microtissues (hMMTs). hMMTs are formed between a pair of opposing microposts in each well, with micropost deflection proportional to the force generated during contraction. In this study, we implement tracer to measure fraction-specific MPS in hMMTs in response to amino acid, ketone and contractile stimulation. We compare stimulation-induced MPS, as measured by both tracers and puromycin incorporation, in hMMTs to that of 2D myotubes derived from the same primary myoblast line. Our findings highlight the enhanced physiological relevance of engineered skeletal muscle compared to 2D myotubes and demonstrate the ability of tracer to capture anabolic responses that are undetected by puromycin-based methods.

## 2. MATERIALS AND METHODS

### 2.1 Human primary myoblast culture and 2D differentiation

A human primary myoblast line was purchased from Cook MyoSite Inc. (SK-1111-P01358-19F; Pittsburgh, PA, USA). The cell line was derived using a proprietary method from a sample of vastus lateralis muscle tissue of a 19-year-old Caucasian female with a BMI of 24 and no known medical conditions. Myoblasts were passaged as previously described (Gulati et al., 2024; Tiper et al., 2025). Briefly, they were plated on collagen-coated tissue culture dishes **(Supplemental Table 1)** in growth medium **(Supplemental Table 1)**, which was replaced every other day. Myoblasts were passaged with trypsin once they reached ∼70 % confluency and replated at a lower density. At passage 8, they were used for 2D myotubes and 3D tissue experiments. The population was ∼95 % CD56^+^ at this passage, as determined by fluorescence-activated cell sorting, indicating a high degree of myogenic purity.

For 2D differentiation, myoblasts were plated at a seeding density of ∼35,000 cells/cm^2^ on collagen-coated 6-well plates **(Supplemental Table 1)**. The cells were cultured in growth medium without added basic fibroblast growth factor for 2 days before transitioning to 2D differentiation medium **(Supplemental Table 1)**. Half the medium was refreshed every other day. Anabolic responses of cultures were assessed on day 5 of differentiation.

### 2.2 MyoTACTIC culture platform fabrication and hMMT seeding

The polydimethylsiloxane (PDMS; Sylgard™ 184 silicone elastomer kit, Dow Corning; Midland, MI, USA) MyoTACTIC 96-well culture platform was fabricated as previously described (Afshar et al., 2020; Lad et al., 2021). After fabrication, the plate was cut into portions containing 4 ± 2 wells. The PDMS portions were sonicated in isopropanol for 20 min, rinsed with dH_2_O, dried in a curing oven at 60-65 °C, and then autoclaved in an instrument sterilization bag. Prior to use, a non-adhesive surface was created by coating the portions with a 5 % Pluronic® F-127 (Cat. #P2443, Sigma-Aldrich; Burlington, MA, USA) solution in dH_2_O overnight at 4 °C. The solution was aspirated prior to hMMT seeding.

hMMTs were seeded in the MyoTACTIC platform as previously described (Afshar et al., 2020; Lad et al., 2021). Primary myoblasts were resuspended in a hydrogel mixture **(Supplemental Table 1)** at a concentration of 11.7 x 10^6^ cells/mL, with thrombin (Cat. #T6884, Sigma-Aldrich) added at 0.37 units/mg of fibrinogen (Cat. #F8630, Sigma-Aldrich) to induce clotting. The resulting hMMTs contained 175,000 cells/tissues. They were cultured in after-seeding growth media **(Supplemental Table 1)** for two days before transitioning to 3D differentiation medium **(Supplemental Table 1)**. Half of the medium was refreshed every other day. Anabolic responses of hMMTs were assessed on day 14 of differentiation.

### 2.3 Anabolic and electrical stimulation conditions

For all conditions, 2D myotubes and 3D hMMTs were first subjected to a 4-hour anabolic knockdown in amino acid-, horse serum- and insulin-free medium **(Supplemental Table 1)**. Following knockdown, the cultures were incubated for 2 hours during the refeeding period in medium supplemented with amino acids, ketones (4 mM β-hydroxybutyrate [Cat. #166898, Sigma-Aldrich) and 1.4 mM lithium acetoacetate [Cat. #A8509, Sigma-Aldrich]), or both amino acids and ketones **(Supplemental Table 1)**. A starvation condition was included as control, where myotubes and hMMTs underwent knockdown without nutrient stimulation. For 3D hMMTs, the effect of contraction on protein synthesis prior to nutrient stimulation was also measured. This involved conditions where hMMTs underwent an electrical stimulation protocol prior to incubation with amino acids and/or ketones. Electrical field stimulation (EFS) was applied to hMMTs between two needle electrodes placed on either side of the microtissue, as previously described (Afshar et al., 2020; Lad et al., 2021). Twitch contractions were elicited using EFS at 1 Hz, 5 V and 60 ms, while tetanic contractions were elicited with EFS at 7 H, 5 V, and 10 ms. The electrical stimulation regimen consisted of 20 twitch contractions followed by 5 tetanus contractions.

### 2.4 Protein synthesis measurements

During the 2-hour refeeding period or the final 2 hours of starvation, *2*50 µM L-*[ring-*^*2*^*H*_*5*_]-phenylalanine (Cat. # DLM-1258-5, Cambridge Isotopes, Tewksbury, MA, USA) was added to the media of each hMMT to assess protein synthesis through tracer incorporation. Additionally, during the last 30 min of feeding or starvation, 1 µM of puromycin (Cat. #PUR333, BioShop; Burlington, ON, Canada) was included in the media of each hMMT to measure protein synthesis via incorporation of puromycin into nascent peptides (Goodman et al., 2011). Immediately before cell lysis, an aliquot of media was collected to measure precursor enrichment for tracer analysis, as described below.

Protein fractions of the homogenate were separated by centrifugation at 1500 x g for 10 min at 4 °C. The supernatant, containing soluble sarcoplasmic proteins, was transferred to a new microtube. The pellet, containing insoluble myofibrillar proteins, was rinsed by adding 500 μL of ice-cold ddH_2_O, centrifuging at 1500 x g for 10 min at 4 °C, and discarding the resulting supernatant (Roberts et al., 2020). Myofibrillar proteins were then solubilized with the addition of 200 μL of 0.3 M NaOH, and heated at 50 °C for 30 min, with vortexing every 10 min. Subsequently, 500 μL of 1 M perchloric acid was added to both the myofibrillar and sarcoplasmic protein tubes to precipitate proteins. The samples were centrifuged at 1600 x g for 20 min at 4 °C, and the supernatant was discarded. The resulting protein pellets were gently rinsed with 70% ice-cold ethanol, centrifuging at 1600 x g for 20 min at 4 °C, and discarding the ethanol supernatant. Proteins were then hydrolyzed into their constituent amino acids by incubation for 48 hours at 110 °C in 750 μL of 0.1 M HCl and 750 μL of Dowex resin (Cat. #Dowex50W-X8–200, Supelco; Bellefonte, PA, USA) in 1 M HCl.

Amino acids were isolated over cation exchange columns constructed from glass wool in 5 mL syringes. The amino acids were eluted with 4 mL of 2 M NH_4_OH, dried down under a steady stream of nitrogen at 80 °C, and resuspended in 200 µL of 0.1 % formic acid. Liquid chromatography tandem mass spectrometry (LC-MS/MS) was then performed to analyze samples, monitoring mass-to-charge ratios (m/z) of 171.1/125 for L-[*ring*-^2^H_5_]Phenylalanine, and 166.1/131 for natural phenylalanine, following established methods for human skeletal muscle (Hannaian et al., 2020).

Fractional synthetic rates (FSR) for both myofibrillar (MyoFSR) and sarcoplasmic (SarcFSR) fractions were calculated as previously described (Atherton et al., 2009) using the equation:

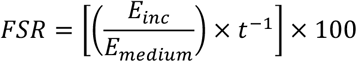

where *E*_*incorp*_ is the enrichment of protein-bound L-[*ring*-^2^H_5_] Phenylalanine, *E*_*medium*_ is the enrichment of the media, and *t* is time (2 hours in all experiments). FSR is expressed in percent per hour.

### 2.5 Immunoblotting

Protein concentrations of the sarcoplasmic fraction were determined by bicinchoninic acid (BCA) assay (Cat. #23227, Thermo Scientific; Waltham, MA, USA). Samples were diluted to equal concentrations, and 10 µg of protein per sample loaded onto 8-16% StainFree Tris-Glycine gels (Cat. <u>#</u>5678105, BioRad; Hercules, CA, USA). The proteins were separated at 120 V for 75 min. Gels were imaged using StainFree Technology (45 sec activation) to quantify total protein content before being transferred onto nitrocellulose membranes at 100 V for 1 hour. The nitrocellulose membranes were blocked for 1 hour at room temperature in 5% skim milk prepared in Tris-buffered saline with 0.1% Tween20™ (TBST). Membranes were then incubated overnight at 4 °C with a primary antibody directed towards puromycin (Cat. #MABE343, RRID: AB_2566826, Sigma-Aldrich) diluted 1:5000 in TBST with 5% bovine serum albumin (BSA). The following day, membranes were washed for 3 x 5min in TBST, and then incubated for 1 hour at room temperature in horseradish peroxidase-conjugated secondary antibody (Cat. #7076, RRID: AB_330924, Cell Signalling Technology; Danvers, MA, USA) diluted 1:5000 in TBST. After another series of 3 x 5 min washes in TBST, membranes were incubated in enhanced chemiluminescence substrate (Cat. #WBKLS0500, Millipore; Burlington, MA, USA ) for 2 min before imaging on a BioRad ChemiDoc system (∼60 sec exposure; Cat. #12003153). Adjusted band intensities was quantified using ImageLab software (RRID:SCR_014210, BioRad), and normalized relative to total protein content as determined by StainFree staining.

### 2.6 Statistical Analysis

In 3D and 2D experiments, tracer and puromycin incorporation were analyzed by pooling 5–7 hMMTs or 3 wells from a 6-well plate, respectively, for each experimental condition. All experiments were performed on three separate occasions, constituting three biological replicates. Values are reported as mean ± standard deviation (SD), and error bars in bar graphs represent SD. Statistical significance was determined using an unpaired, two-tailed *t*-test for comparison between two groups, and a one-way ANOVA followed by Tukey’s post hoc test for multiple group comparisons. Statistical significance was set to *P* ≤ 0.05 for all tests. Details of pooled tissue counts, means ±SD, statistical tests, and P-values for measurements of protein synthesis are provided in **Supplemental Table 2** (2D studies) and **Supplemental Table 3** (3D studies). Simple linear regression was performed to examine the relationships between variables. The x- and y-axis metrics, F-Statistic, P-value, slope, and coefficient of determination for each regression are provided in **Supplemental Table 4**. All graphs were generated, and statistical analyses were performed using GraphPad Prism 10.0 (GraphPad Software, RRID:SCR_002789; Boston, MA, USA).

## 3. RESULTS

### 3.1 Tracer provides greater sensitivity for measuring MPS than puromycin

To compare measurements of MPS derived from stable isotope tracers and puromycin, we assessed L-[ring-^2^H_5_]-phenylalanine and puromycin incorporation during acute stimulation of 2D myotubes with amino acids, ketones (β-hydroxybutyrate and lithium acetoacetate), or both, following anabolic knockdown (n = 3 wells per condition across N = 3 independent experiments; **Figure 1A**). MPS in starved myotubes during knockdown served as the baseline. We found sarcoplasmic FSR increased significantly with amino acid treatment, whereas ketone stimulation had no effect (*p* = 0.002 and *p* > 0.999, respectively; one-way ANOVA; **Figure 1B**). Combined amino acid and ketone treatment significantly increased FSR relative to starved myotubes but was not significantly different from amino acids alone (*p* = 0.0004 and *p* = 0.481, respectively; one-way ANOVA). A similar trend was observed for the myofibrillar FSR, which increased significantly with amino acid stimulation but remined unaffected by ketones (*p* = 0.002 and *p* = 0.996, respectively; one-way ANOVA; **Figure 1C**). Combined amino acid and ketone treatment significantly increased myofibrillar FSR relative to starved myotubes but again did not differ significantly from amino acids alone (*p* = 0.0008 and *p* = 0.863, respectively; one-way ANOVA). Linear regression analysis revealed a very strong and statistically significant positive relationship between myofibrillar and sarcoplasmic FSRs, with higher synthesis rates observed in the sarcoplasmic fraction across stimulation conditions (R^2^ = 0.990, m = 0.700, *p* < 0.001; **Figure 1D**). In contrast, puromycin-derived measurements of MPS failed to capture nutrient-induced responses in myotubes **(Figure 1E, F)**. Puromycin incorporation did not increase significantly with amino acid, ketone, or a combined nutrient stimulation (*p* = 0.915, p = 0.971, and *p* = 0.945, respectively; one-way ANOVA). This is consistent with a linear regression showing minimal correlation between puromycin incorporation and sarcoplasmic FSR (R^2^ = 0.0337, m = 0.0727, *p* 287 = 0.568; **Figure 1G**). These results demonstrate the greater sensitivity conferred by a tracer-based approach for quantifying MPS at longer timescales, as nutrient-stimulated increases in protein synthesis were detected using stable phenylalanine isotopes, which puromycin failed to capture.

**Figure 1.**
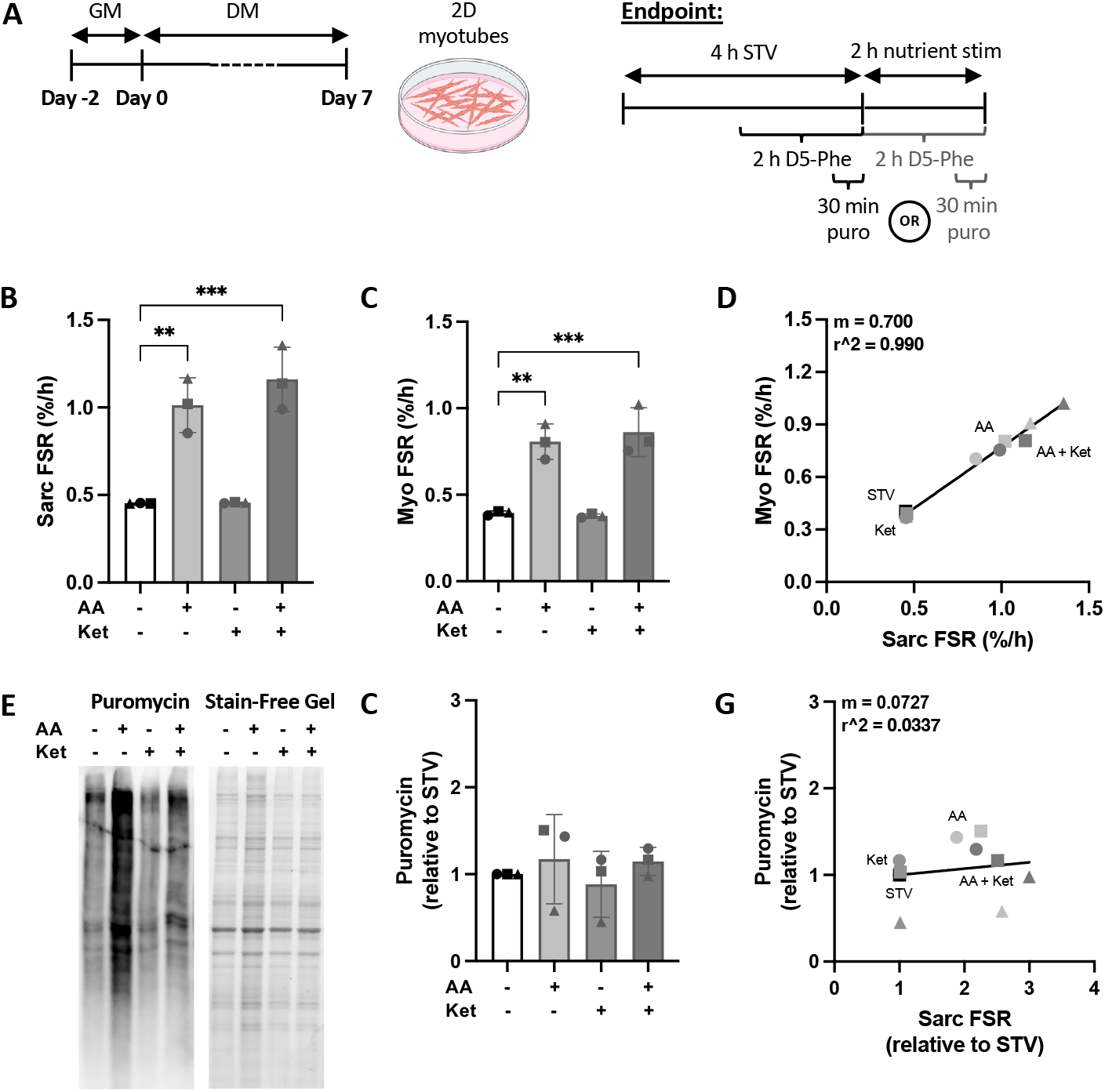
Stable isotope-versus puromycin-derived measurements of nutrient stimulated muscle protein synthesis (MPS) in 2D myotubes. (A) Schematic overview of the timeline for 2D myotube culture and endpoint MPS measurements. Cells were cultured in growth media (GM) for 2 days, followed by differentiation media (DM) for 5 days. MPS was measured via incorporation of L-*[ring-*^*2*^*H*_*5*_]-phenylalanine (D5-Phe) and puromycin during anabolic knockdown (starved [STV]), or stimulation with amino acids (AA), ketones (Ket), or both (AA + Ket) following knockdown. Sarcoplasmic (sarc; B) and myofibrillar (myo; C) fractional synthetic rates (FSR) measured via D5-Phe incorporation into sarcoplasmic and myofibrillar proteins, respectively. (D) Linear regression modelling the relationship between sarcoplasmic and myofibrillar FSR. (E) Quantification of puromycin-labeled peptides normalized to STV. (F) Representative image of western blot analysis for puromycin alongside the corresponding stain-free total protein image. (G) Linear regression modelling the relationship between sarcoplasmic FSR and puromycin incorporation. n = 3 wells pooled per condition across N = 3 independent experiments. Graphs display means ±SD. Circular, triangular, and square dots represent values from the first, second and third biological replicate, respectively. Statistical significance was determined using one-way ANOVA followed by Tukey’s post hoc test; significance is only displayed for conditions relative to STV; ** *p* ≤ 0.01, *** *p* ≤ 0.001.

### 3.2 Ketones and contraction do not enhance amino acid stimulated protein synthesis in hMMTs

To further compare the sensitivity of tracer- and puromycin-based approaches for quantifying MPS and to investigate the combined effect of contractile and nutrient stimulation on MPS, we repeated the experiment in hMMTs with the inclusion of a contraction regimen **(Figure 2A)**. L-[ring-^2^H_5_]-phenylalanine and puromycin incorporation were assessed during acute stimulation of hMMTs with amino acids, ketones, or both, following anabolic knockdown with or without an accompanying electrical contraction protocol prior to treatment. Consistent with the results in 2D myotubes, we found sarcoplasmic FSR increased significantly with amino acid treatment, whereas ketone stimulation had no effect (*p* < 0.0001 and *p* = 0.999, respectively; one-way ANOVA; **Figure 2B**). Combined amino acid and ketone treatment significantly increased FSR relative to starved myotubes but was not significantly different from amino acids alone (*p* < 0.0001 and *p* = 0.919, respectively; one-way ANOVA). Additionally, contractile stimulation prior to amino acid treatment significantly increased FSR relative to starved myotubes but did not enhance sarcoplasmic FSR beyond the level achieved with only amino acids (*p* = < 0.0001 and *p* = 0.998, respectively; one-way ANOVA). Increases in myofibrillar FSR in response to amino acid stimulation, either alone, or in combination with ketones, contraction, or both, were not significant (*p* = 0.159, *p* = 0.173, *p* = 0.198, and *p* = 0.100, respectively; one-way ANOVA; **Figure 2C**). Despite the myofibrillar fraction being less responsive to anabolic stimuli, linear regression analysis revealed a strong and statistically significant positive relationship between myofibrillar and sarcoplasmic FSRs (R^2^ = 0.807, m = 0.313, *p* < 0.0001; **Figure 2D**). The Puromycin-based approach for quantifying MPS did not capture the responses of hMMTs to the anabolic stimuli and produced data with large variability (means **±** SD and p-values are presented in **Supplemental Table 3**; one-way ANOVA; **Figure 2E, F)**. This aligns with a linear regression showing a weak and statistically insignificant relationship between puromycin incorporation and sarcoplasmic FSR (R^2^ 314 = 0.123, m = -0.200, *p* = 0.120; **Figure 2G**). Thus, using stable isotope tracers to measure MPS, we observed that acute amino acid-stimulated protein synthesis in hMMTs is not enhanced by prior contractile stimulation or combined ketone treatment. There were no differences in puromycin-derived measurements of MPS, further emphasizing the superiority of the tracer-based approach for queries requiring longer pulse lengths to deconvolute distinctions.

**Figure 2.**
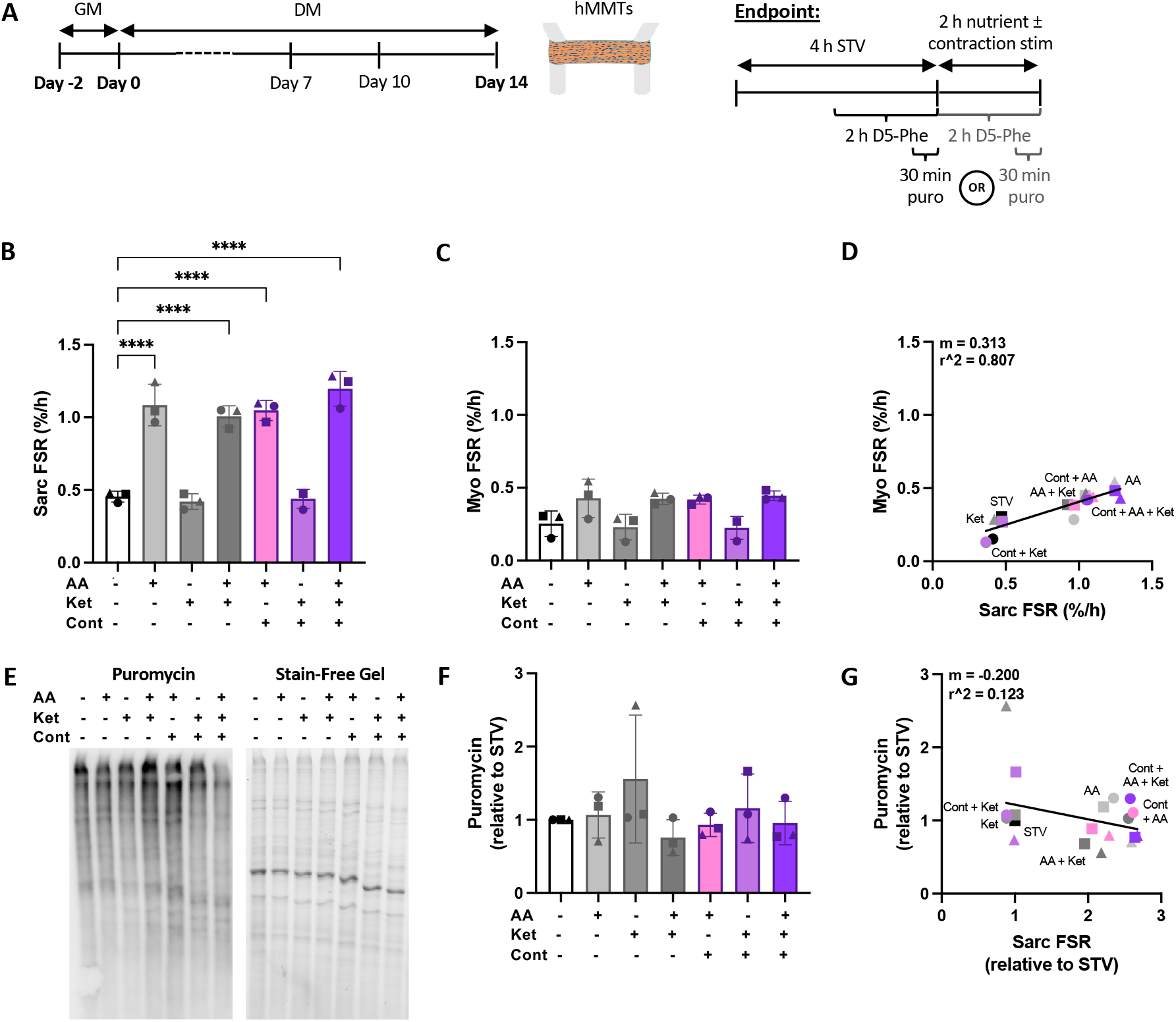
Stable isotope-versus puromycin-derived measurements of nutrient and contraction stimulated muscle protein synthesis (MPS) in human skeletal muscle microtissues (hMMTs). (A) Schematic overview of the timeline for hMMT culture and endpoint MPS measurements. hMMTs were cultured in after-seeding growth media (GM) for 2 days, followed by differentiation media for 14 days. MPS was measured via incorporation of L-*[ring-*^*2*^*H*_*5*_]-phenylalanine (D5-Phe) and puromycin during anabolic knockdown (starved [STV]), or stimulation with amino acids (AA), ketones (Ket), or both (AA + Ket) following knockdown, with or without an accompanying electrical contraction (Cont) regime prior to treatment. Sarcoplasmic (sarc; B) and myofibrillar (myo; C) fractional synthetic rates (FSR) measured via D5-Phe incorporation into sarcoplasmic and myofibrillar proteins, respectively. (D) Linear regression modelling the relationship between sarcoplasmic and myofibrillar FSR. (E) Quantification of puromycin-labeled peptides normalized to STV. (F) Representative image of western blot analysis for puromycin alongside the corresponding stain-free total protein image. (G) Linear regression modelling the relationship between sarcoplasmic FSR and puromycin incorporation. n = 5-7 hMMTs pooled per condition across N = 3 independent experiments. Graphs display means ±SD. Circular, triangular and square dots represent values from the first, second and third biological replicate, respectively. Statistical significance was determined using one-way ANOVA followed by Tukey’s post hoc test; significance is only displayed for conditions relative to STV; **** *p* ≤ 0.0001.

### 3.3 Myofibrillar protein fraction of hMMTs exhibits greater stability than in 2D myotubes

We performed linear regression analyses to assess the relationships between sarcoplasmic FSR **(Figure 3A)**, myofibrillar FSR **(Figure 3B)**, and puromycin incorporation **(Figure 3C)** in 2D myotubes and 3D hMMTs under nutrient stimulation. Both sarcoplasmic FSR and myofibrillar FSR were highly correlated across the two models (R^2^ = 0.935 and R^2^ = 0.986, respectively). The sarcoplasmic FSR regression slope (m = 0.922, *p* = 0.033) suggests similar nutrient-stimulated responses in the sarcoplasmic protein fraction of myotubes and hMMTs, whereas the myofibrillar FSR regression slope (m = 0.409, *p* = 0.007) indicates reduced responsiveness of the myofibrillar fraction of hMMTs. Accordingly, the sarcoplasmic-to-myofibrillar FSR ratio across nutrient stimulation conditions was higher in hMMTs (2.132 ± 0.378) compared to myotubes (1.239 ± 0.085; *p* = 0.0036; **Table 1**). Although puromycin incorporation showed moderate correlation (R^2^ = 0.638), its regression slope was non-significant (*p* = 0.201). Comparison of tracer-based measurement of MPS in 2D myotubes and hMMTs reflects the distinct biological properties of these skeletal muscle models, with the myofibrillar fraction in hMMTs exhibiting greater stability than that in myotubes.

**Table 1.** Sarcoplasmic-to-myofibrillar fractional synthetic rate (FSR) ratio across stimulation conditions in 2D myotubes and 3D hMMTs.

| Culture | Stimulation condition | Sarcoplasmic-to-myofibrillar FSR ratio | Mean ratio ± SD | P value |
| --- | --- | --- | --- | --- |
| 2D myotubes | STV | 1.15 | 1.239 ± 0.085 | *p* = 0.0036 |
|  | AA | 1.26 |  |  |
|  | Ket | 1.21 |  |  |
|  | AA + Ket | 1.35 |  |  |
| 3D hMMTs | STV | 1.79 | 2.132 ± 0.378 | |
|  | AA | 2.54 |  |  |
|  | Ket | 1.83 |  |  |
|  | AA + Ket | 2.37 |  |  |
|  | Cont + AA | 2.50 | Contractile stimulation conditions not included |  |
|  | Cont + Ket | 1.96 |  |  |
|  | Cont + AA + Ket | 2.69 |  |  |

Abbreviations: STV = starved, AA = amino acid, Ket = ketone.

**Figure 3.**
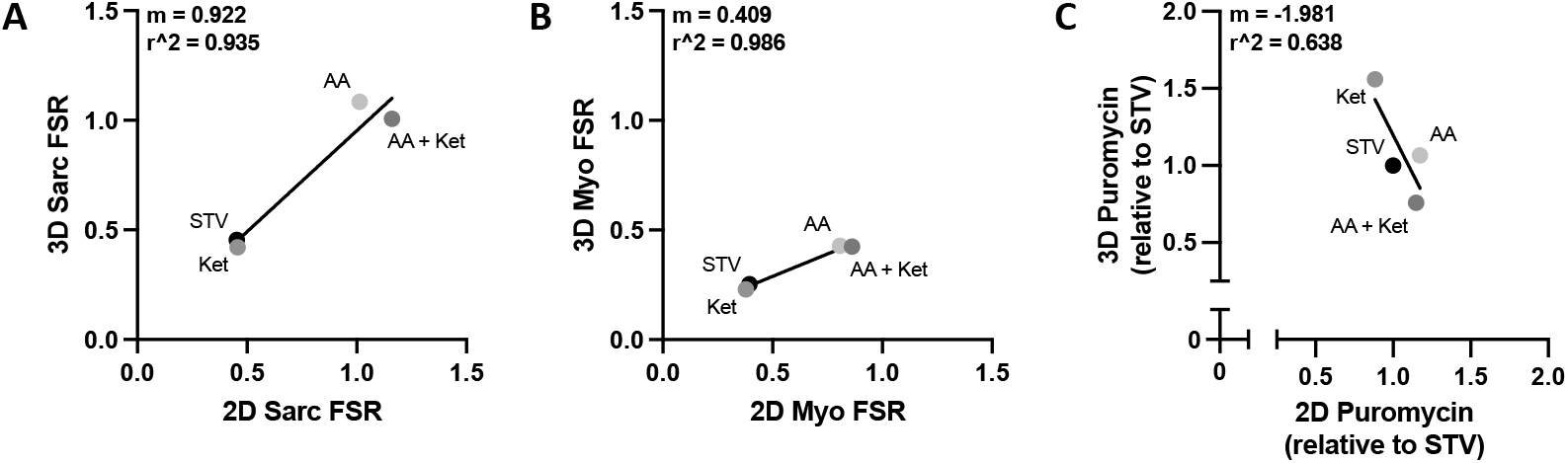
Comparison of stable isotope- and puromycin-derived measurements of nutrient stimulated protein synthesis in 2D myotubes versus 3D human skeletal muscle microtissues (hMMTs). Linear regressions illustrating the relationships for measurements of sarcoplasmic fractional synthetic rate (Sarc FSR; A), myofibrillar fractional synthetic rate (Myo FSR; B), and puromycin incorporation (C) in 2D myotubes versus 3D hMMTs. Measurements for starved (STV), amino acid (AA), ketone (Ket), and combined amino acid and ketone (AA + Ket) stimulation conditions were used in the analysis. Data points represent the mean of N = 3 biological replicates.

## 4. DISCUSION

In the present study, we demonstrate that stable isotope amino acid tracer was more sensitive to detecting acute nutrient-induced changes in protein synthesis than puromycin incorporation. Stable isotopes enabled fraction-specific measurements of protein synthesis in skeletal muscle *in vitro* and captured anabolic responses that puromycin failed to detect. Although there was a high level of agreement between the responses of 2D and 3D skeletal muscle models, 3D hMMTs displayed a sarcoplasmic-to-myofibrillar FSR ratio of ∼2:1 across conditions **(Figure 2)**, compared to a ratio of ∼1:1 in 2D myotubes **(Figure 1)**. This finding can be reinforced in future studies by evaluating multiple donor cell lines. Notably, the ratio ∼2:1 observed for hMMTs closely resembles that of native human skeletal muscle (Abou Sawan et al., 2021; Moore, Tang, et al., 2009), demonstrating the ability of bioengineered models to better recapitulate the fraction-specific MPS responses characteristic of human muscle. However, the ketones β-hydroxybutyrate and lithium acetoacetate did not enhance muscle protein synthesis, either in the absence of amino acids–likely due to the lack of a basal amino acid pool found *in vivo*–or in the presence of supraphysiological concentrations of essential amino acids. Furthermore, contractile stimulation did not potentiate the response to amino acid sufficiency during the acute post-stimulation period, suggesting that a longer duration may be required to observe an effect. Overall, our results highlight the value of the tracer-based approach in providing insights into protein synthesis not revealed by puromycin.

### 4.1 Limitations of puromycin-based approach likely explain lack of observed differences

Puromycin incorporation is a convenient and cost-effective method to measure nascent peptide synthesis, but it has several limitations that affect its utility. Unlike stable isotope tracers, puromycin is neither chemically nor functionally identical to naturally occurring amino acids. As a potent inhibitor of translation, puromycin terminates protein synthesis upon incorporation into elongating peptide chains, making its integration into functional proteins highly unlikely. Furthermore, puromycin assays are performed by immunoblotting the soluble sarcoplasmic fraction of cell lysates. Although this approach only assesses the sarcoplasmic fraction, it is more reflective of a ‘mixed’ protein fraction, because puromycin-tagged truncated proteins often fail to reach their intended cellular compartments and remain in the sarcoplasm. The inherently lower precision of immunoblotting, when compared to LC-MS/MS (Aebersold et al., 2013), likely contributes to the greater variability observed between biological replicates in our puromycin-based measurements. Additionally, puromycin labelling does not accurately measure mRNA translation rates in energetically challenged cells. In glucose- and amino acid-starved HEK293 cells, puromycin incorporation showed a significantly smaller reduction in protein synthesis compared to radiolabelled amino acid ([^35^S]methionine) incorporation (Marciano et al., 2018). This suggests that puromycin may underestimate the extent of translational inhibition under nutrient-deprived conditions, such as in the 2D myotubes and hMMTs in our study following anabolic knockdown. Another key limitation is that the shorter 30-minute incorporation period used for the puromycin, compared to the 2 hours for tracer, may capture distinct protein synthesis trends. Short incorporation times preferentially bias measurement towards proteins with rapid turnover rates (Miller et al., 2015), thereby underrepresenting proteins with slower turnover dynamics. These limitations highlight the need for cautious interpretation of puromycin-derived measurements of protein synthesis.

### 4.2 Absence of ketone-induced increase in MPS likely due to insufficient amino acid availability

Ketone supplementation with β-hydroxybutyrate and lithium acetoacetate in amino acid-free DMEM failed to stimulate protein synthesis in 2D myotubes or hMMTs relative to control amino acid-free DMEM following anabolic knockdown. In contrast, ketone monoester supplementation in fasted human skeletal muscle increased MPS to a level comparable to a small bolus (10g) of whey protein (Hannaian et al., 2024). This disparity suggests that while ketones can stimulate MPS, they require a basal concentration of amino acids as substrates for protein translation. *In vivo*, human plasma in the overnight fasted (post-absorptive) state maintains amino acids levels sufficient to support protein synthesis, whereas *in vitro* models lack this physiological baseline. Moreover, our studies demonstrated that ketones did not enhance protein synthesis in the presence of supraphysiological amino acid concentrations found in DMEM in either 2D myotubes or hMMTs. This finding aligns with evidence from human skeletal muscle, where ketone monoester supplementation did not exert additive effects when combined with whey protein ingestion (Hannaian et al., 2024). Future work should investigate ketone supplementation in media containing physiological concentrations of amino acids that more closely resemble human serum composition.

### 4.3 Contraction-mediated increases in MPS likely missed due to short measurement duration

While amino acid provision increased MPS in hMMTs, contraction did not provide any additional enhancement. This differs from *in vivo* human data, where resistance exercise (i.e., contraction) combined with protein ingestion induces a greater stimulation of MPS than protein alone, primarily due to elevated myofibrillar protein synthesis (Burd et al., 2011; Moore, Tang, et al., 2009). One study reported that myofibrillar protein synthesis was significantly higher in the exercise-plus-fed condition compared to feeding alone at 3- and 5-hours post-exercise (Moore, Tang, et al., 2009). Given that our labeling period was limited to 2 hours post-stimulation, the full potentiation effect of contraction may not have been captured. To address this, future studies should extend the MPS measurement window and include a contraction-only control. This would establish whether hMMTs exhibit increased myofibrillar MPS in response to the synergistic effect of contraction and amino acid provision, but not contraction alone, as is observed *in vivo*.

## 5. Conclusions

Overall, we demonstrate the utility of stable isotope tracers for measuring protein synthesis in 2D and 3D human skeletal muscle culture models. Our findings indicate hMMTs generated in the MyoTACTIC platform better recapitulate human physiological responses to nutrient stimulation compared to 2D myotubes derived from the same primary cell line. Future work in this space should prioritize use of physiologically relevant skeletal muscle models and utilize stable isotope-derived measurements to enhance the accuracy of muscle protein synthesis assessments.

## AUTHOR CONTRIBUTIONS

Cassidy T Tinline-Goodfellow and Yekaterina Tiper were responsible for the acquisition and analysis of the data presented in this work. All authors contributed to the study’s conception and design, and data interpretation. Cassidy T Tinline-Goodfellow and Yekaterina Tiper drafted the manuscript. All authors revised it critically for important intellectual content. The final version of the manuscript has been approved by all authors, who agree to be accountable for all aspects of the work in ensuring that questions related to the accuracy or integrity of any part of the work are appropriately investigated and resolved. All persons designated as authors qualify for authorship, and all those who qualify for authorship are listed.

## FUNDING

This project was supported by a University of Toronto XSeed Award to Daniel Moore and Penney Gilbert and a Canada First Research Excellence Fund “Medicine by Design (MbD)” grant awarded to Penney M Gilbert. Yekaterina Tiper received a Wildcat Graduate Scholarship and Doctoral Completion Award from the Department of Biomedical Engineering. Penney M Gilbert also holds a Canada Research Chair in Endogenous Repair award (#950-231201).

## CONFLICT OF INTEREST

No conflicts of interest, financial, or otherwise, are declared by the authors.

## DATA AVAILABILITY STATEMENT

The data that support the findings of this study are available from the corresponding authors upon reasonable request.

## Supplemental Tables

**Supplemental Table 1.** Composition of media and solutions used for human primary myoblast expansion, 2D myotube differentiation, hMMT culture, and endpoint anabolic stimulation.

| Media/solution | Composition |
| --- | --- |
| Rat tail collagen coating solution for myoblast passaging | ~16 µg/mL rat tail Collagen I (Cat. #A1048301, Gibco; Waltham, MA, USA ) in 0.1 M acetic acid solution prepared with dH2O |
| Myoblast growth medium | Ham's F-10 nutrient mix (Cat. #318-050-CL, Wisent Bioproducts; Saint-Jean Baptiste, QC, Canada), 20% fetal bovine serum (FBS; Cat. #12483020, Gibco), 5 ng/mL basic fibroblast growth factor (bFGF; Cat. #11343625, ImmunoTools; Friesoythe, Germany), 1% Penicillin-Streptomycin (P/S; Cat. #15140122, Gibco) |
| Rat tail collagen coating solution for 2D differentiation | ~50 µg/mL rat tail Collagen I in 0.1 M acetic acid solution prepared with dH2O |
| 2D differentiation medium | High glucose (4.5 g/L) Dulbecco's Modified Eagle's Medium (DMEM; Cat. #11995065, Gibco), 2% horse serum (HS; Cat. #16050114, Gibco), 10 µg/mL insulin (Cat. #I6634, Sigma-Aldrich), 1% P/S |
| Fibrinogen stock solution | 10 mg/mL fibrinogen in 0.9% (wt/v) NaCl solution prepared with dH2O |
| Hydrogel master mix | High glucose DMEM (40% v/v), 4 mg/mL fibrinogen (40% v/v), Geltrex™ (20% v/v; Cat. #A1413202, Gibco) |
| After-seeding growth medium | Ham's F-10 nutrient mix, 20% FBS, 1.5 mg/mL 6-aminocaproic acid (ACA; 3% v/v; Cat. #A2504, Sigma-Aldrich), 1% P/S |
| 3D differentiation medium | High glucose DMEM, 2% HS, 2 mg/mL ACA (4% v/v), 10 µg/mL insulin, 1% P/S |
| Anabolic knockdown medium | Low glucose (1.0 g/L) DMEM w/o amino acids (Cat. #D9800-13, United States Biological Corporation; Salem, MA, USA), 3.5 g/L D-glucose (bringing total glucose up to 4.5 g/L), 1 mM sodium pyruvate (Cat. 11360070, Gibco), 3.7 g/L sodium bicarbonate, 2 mg/mL ACA (4% v/v), 1% P/S |
| Amino acid stimulation medium | High glucose DMEM, 2 mg/mL ACA (4% v/v), 1% P/S |
| Ketone stimulation medium | Low glucose DMEM with w/o amino acids, 3.5 g/L D-glucose, 1 mM sodium pyruvate, 3.7 g/L sodium bicarbonate, 4 mM 3-hydroxybutyrate, 1.4 mM lithium acetoacetate, 2 mg/mL ACA (4% v/v), 1% P/S |
| Amino acid and ketone stimulation medium | High glucose DMEM, 4 mM 3-hydroxybutyrate, 1.4 mM lithium acetoacetate, 2 mg/mL ACA (4% v/v), 1% P/S |

**Supplemental Table 2.**
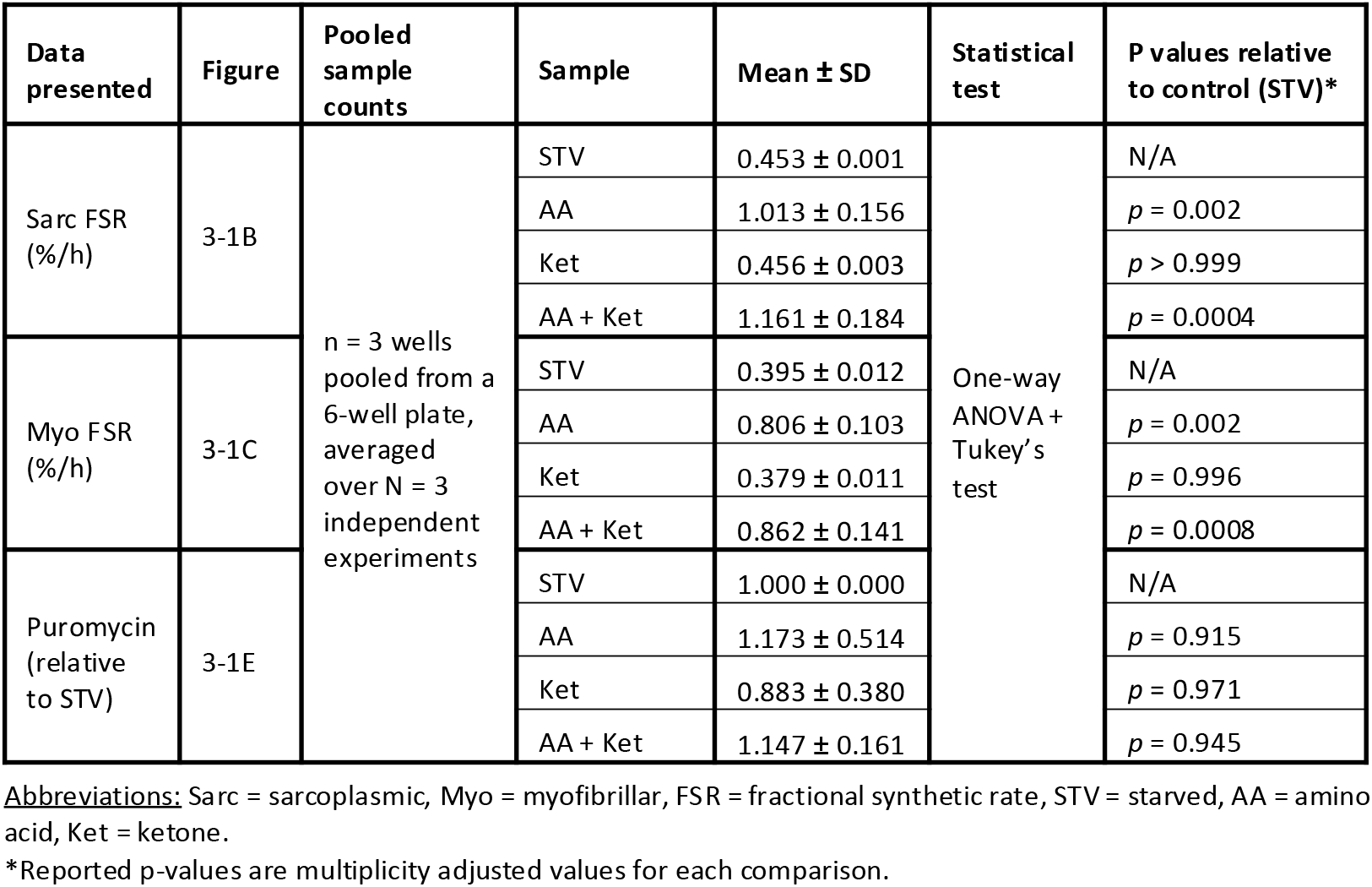
Pooled sample counts, means ± standard deviation (SD), statistical tests, and significance values for measurements of protein synthesis in 2D studies.

**Supplemental Table 3.** Pooled sample counts, means ± standard deviation (SD), statistical tests, and significance values for measurements of protein synthesis in 3D studies.

| Data presented | Figure | Pooled sample counts | Sample | Mean ± SD | Statistical test | P values relative to control (STV)* |
| --- | --- | --- | --- | --- | --- | --- |
| Sarc FSR (%/h) | 3-2B | n = 5-7 pooled hMMTs, averaged over N = 3 independent experiments | STV | 0.454 ± 0.037 | One-way ANOVA + Tukey's test | N/A |
| | | | AA | 1.085 ± 0.143 | | $p < 0.0001$ |
| | | | Ket | 0.420 ± 0.054 | | $p = 0.999$ |
| | | | AA + Ket | 1.006 ± 0.073 | | $p < 0.0001$ |
| | | | Cont + AA | 1.048 ± 0.069 | | $p < 0.0001$ |
| | | | Cont + Ket | 0.439 ± 0.065 | | $p > 0.999$ |
| | | | Cont + AA + Ket | 1.197 ± 0.120 | | $p < 0.0001$ |
| Myo FSR (%/h) | 3-2C | | STV | 0.253 ± 0.087 | | N/A |
| | | | AA | 0.428 ± 0.132 | | $p = 0.159$ |
| | | | Ket | 0.230 ± 0.089 | | $p > 0.999$ |
| | | | AA + Ket | 0.424 ± 0.039 | | $p = 0.173$ |
| | | | Cont + AA | 0.419 ± 0.031 | | $p = 0.198$ |
| | | | Cont + Ket | 0.224 ± 0.079 | | $p = 0.999$ |
| | | | Cont + AA + Ket | 0.445 ± 0.033 | | $p = 0.100$ |
| Puromycin (relative to STV) | 3-2E | | STV | 1.000 ± 0.000 | | N/A |
| | | | AA | 1.066 ± 0.316 | | $p > 0.999$ |
| | | | Ket | 1.558 ± 0.874 | | $p = 0.677$ |
| | | | AA + Ket | 0.758 ± 0.243 | | $p = 0.990$ |
| | | | Cont + AA | 0.931 ± 0.160 | | $p > 0.999$ |
| | | | Cont + Ket | 1.158 ± 0.469 | | $p = 0.999$ |
| | | | Cont + AA + Ket | 0.955 ± 0.297 | | $p > 0.999$ |

**Abbreviations:** Sarc = sarcoplasmic, Myo = myofibrillar, FSR = fractional synthetic rate, STV = starved, AA = amino acid, Ket = ketone, Cont = contraction.
\*Reported p-values are multiplicity adjusted values for each comparison.

**Supplemental Table 4.** The x- and y-axis metrics, F-statistic, significance value, slope, and coefficient of determination for each linear regression model.

| Metric |  | Figure | F-Statistic | P-Value | Slope | R<sup>2</sup> |
| --- | --- | --- | --- | --- | --- | --- |
| X-axis | Y-axis |  |  |  |  |  |
| 2D Sarc FSR | 2D Myo FSR | 3-1D | F(1,10) = 979.2 | $p < 0.001$ | m = 0.700 | R<sup>2</sup> = 0.990 |
| 2D Sarc FSR | 2D Puromycin | 3-1G | F(1,10) = 0.349 | $p = 0.568$ | m = 0.0727 | R<sup>2</sup> = 0.0337 |
| 3D Sarc FSR | 3D Myo FSR | 3-2D | F(1,19) = 79.67 | $p < 0.0001$ | m = 0.313 | R<sup>2</sup> = 0.807 |
| 3D Sarc FSR | 3D Puromycin | 3-2G | F(1,19) = 2.653 | $p = 0.120$ | m = -0.200 | R<sup>2</sup> = 0.123 |
| 2D Sarc FSR | 3D Sarc FSR | 3-3A | F(1,2) = 28.75 | $p = 0.033$ | m = 0.922 | R<sup>2</sup> = 0.935 |
| 2D Myo FSR | 3D Myo FSR | 3-3B | F(1,2) = 138.7 | $p = 0.007$ | m = 0.409 | R<sup>2</sup> = 0.986 |
| 2D Puromycin | 3D Puromycin | 3-3C | F(1,2) = 3.521 | $p = 0.201$ | m = -1.981 | R<sup>2</sup> = 0.638 |

**Abbreviations:** Sarc = sarcoplasmic, Myo = myofibrillar, FSR = fractional synthetic rate.

## Notes

### Competing Interest Statement

The authors have declared no competing interest.

## References

Abou Sawan, S., Hodson, N., Tinline-Goodfellow, C., West, D. W. D., Malowany, J. M., Kumbhare, D., & Moore, D. R. (2021). Incorporation of Dietary Amino Acids into Myofibrillar and Sarcoplasmic Proteins in Free-Living Adults Is Influenced by Sex, Resistance Exercise, and Training Status. The Journal of Nutrition, 151(11), 3350–3360. 10.1093/jn/nxab261

Aebersold, R., Burlingame, A. L., & Bradshaw, R. A. (2013). Western blots versus selected reaction monitoring assays: Time to turn the tables? Molecular & Cellular Proteomics, 12(9), 2381–2382. 10.1074/mcp.E113.031658

Afshar, M. E., Abraha, H. Y., Bakooshli, M. A., Davoudi, S., Thavandiran, N., Tung, K., Ahn, H., Ginsberg, H. J., Zandstra, P. W., & Gilbert, P. M. (2020). A 96-well culture platform enables longitudinal analyses of engineered human skeletal muscle microtissue strength. Scientific Reports, 10(1), 6918. 10.1038/s41598-020-62837-8

Atherton, P. J., Szewczyk, N. J., Selby, A., Rankin, D., Hillier, K., Smith, K., Rennie, M. J., & Loughna, P. T. (2009). Cyclic stretch reduces myofibrillar protein synthesis despite increases in FAK and anabolic signalling in L6 cells. The Journal of Physiology, 587(14), 3719–3727. 10.1113/jphysiol.2009.169854

Burd, N. A., West, D. W. D., Moore, D. R., Atherton, P. J., Staples, A. W., Prior, T., Tang, J. E., Rennie, M. J., Baker, S. K., & Phillips, S. M. (2011). Enhanced amino acid sensitivity of myofibrillar protein synthesis persists for up to 24 h after resistance exercise in young men. The Journal of Nutrition, 141(4), 568–573. 10.3945/jn.110.135038

Crossland, H., Smith, K., Atherton, P. J., & Wilkinson, D. J. (2020). A novel stable isotope tracer method to simultaneously quantify skeletal muscle protein synthesis and breakdown. Metabolism Open, 5, 100022. 10.1016/j.metop.2020.100022

Goodman, C. A., Mabrey, D. M., Frey, J. W., Miu, M. H., Schmidt, E. K., Pierre, P., & Hornberger, T. A. (2011). Novel insights into the regulation of skeletal muscle protein synthesis as revealed by a new nonradioactive in vivo technique. The FASEB Journal, 25(3), 1028–1039. 10.1096/á.10-168799

Gran, P., & Cameron-Smith, D. (2011). The actions of exogenous leucine on mTOR signalling and amino acid transporters in human myotubes. BMC Physiology, 11(1), 10. 10.1186/1472-6793-11-10

Gulati, N., Davoudi, S., Xu, B., Rjaibi, S. T., Jacques, E., Pham, J., Fard, A., McGuigan, A. P., & Gilbert, P. M. (2024). Mini-MEndR: a miniaturized 96-well predictive assay to evaluate muscle stem cell-mediated repair. BMC Methods, 1(1), 5. 10.1186/s44330-024-00005-4

Hannaian, S. J., Hodson, N., Sawan, S. A., Mazzulla, M., Kato, H., Matsunaga, K., Waskiw-Ford, M., Duncan, J., Kumbhare, D. A., & Moore, D. R. (2020). Leucine-enriched amino acids maintain peripheral mTOR-Rheb localization independent of myofibrillar protein synthesis and mTORC1 signaling postexercise. Journal of Applied Physiology, 129(1), 133–143. 10.1152/japplphysiol.00241.2020

Hannaian, S. J., Lov, J., Hawley, S. E., Dargegen, M., Malenda, D., Gritsas, A., Gouspillou, G., Morais, J. A., & Churchward-Venne, T. A. (2024). Acute ingestion of a ketone monoester, whey protein, or their co-ingestion in the overnight postabsorptive state elicit a similar stimulation of myofibrillar protein synthesis rates in young males: a double-blind randomized trial. The American Journal of Clinical Nutrition, 119(3), 716–729. 10.1016/j.ajcnut.2024.01.004

Hofemeier, A. D., Limon, T., Muenker, T. M., Wallmeyer, B., Jurado, A., Afshar, M. E., Ebrahimi, M., Tsukanov, R., Oleksiievets, N., Enderlein, J., Gilbert, P. M., & Betz, T. (2021). Global and local tension measurements in biomimetic skeletal muscle tissues reveals early mechanical homeostasis. ELife, 10, e60145. 10.7554/eLife.60145

Huang, Z., Chen, D., Zhang, K., Yu, B., Chen, X., & Meng, J. (2007). Regulation of myostatin signaling by c-Jun N-terminal kinase in C2C12 cells. Cellular Signalling, 19(11), 2286–2295. 10.1016/j.cellsig.2007.07.002

Khodabukus, A. (2021). Tissue-Engineered Skeletal Muscle Models to Study Muscle Function, Plasticity, and Disease. Frontiers in Physiology, 12, 619710. 10.3389/fphys.2021.619710

Kim, I. Y., Suh, S. H., Lee, I. K., & Wolfe, R. R. (2016). Applications of stable, nonradioactive isotope tracers in in vivo human metabolic research. Experimental and Molecular Medicine, 48(1), e203. 10.1038/emm.2015.97

Lad, H., Musgrave, B., Ebrahimi, M., & Gilbert, P. M. (2021). Assessing Functional Metrics of Skeletal Muscle Health in Human Skeletal Muscle Microtissues. Journal of Visualized Experiments: JoVE, 2021(168), e62307. 10.3791/62307

Madden, L., Juhas, M., Kraus, W. E., Truskey, G. A., & Bursac, N. (2015). Bioengineered human myobundles mimic clinical responses of skeletal muscle to drugs. ELife, 4, e04885. 10.7554/eLife.04885

Marciano, R., Leprivier, G., & Rotblat, B. (2018). Puromycin labeling does not allow protein synthesis to be measured in energy-starved cells correspondence. Cell Death and Disease, 9(2), 39. 10.1038/s41419-017-0056-x

Merz, K. E., & Thurmond, D. C. (2020). Role of skeletal muscle in insulin resistance and glucose uptake. Comprehensive Physiology, 10(3), 785–809. 10.1002/cphy.c190029

Miller, B. F., Wolff, C. A., Iii, F. F. P., Shipman, P. D., & Hamilton, K. L. (2015). Modeling the contribution of individual proteins to mixed skeletal muscle protein synthetic rates over increasing periods of label incorporation. J Appl Physiol, 118, 655–661. 10.1152/japplphysiol.00987.2014.-Advances

Moore, D. R. (2019). Maximizing Post-exercise Anabolism: The Case for Relative Protein Intakes. Frontiers in Nutrition, 6, 147. 10.3389/fnut.2019.00147

Moore, D. R., Robinson, M. J., Fry, J. L., Tang, J. E., Glover, E. I., Wilkinson, S. B., Prior, T., Tarnopolsky, M. A., & Phillips, S. M. (2009). Ingested protein dose response of muscle and albumin protein synthesis after resistance exercise in young men. The American Journal of Clinical Nutrition, 89(1), 161–168. 10.3945/ajcn.2008.26401

Moore, D. R., Tang, J. E., Burd, N. A., Rerecich, T., Tarnopolsky, M. A., & Phillips, S. M. (2009). Differential stimulation of myofibrillar and sarcoplasmic protein synthesis with protein ingestion at rest and after resistance exercise. The Journal of Physiology, 587(4), 897–904. 10.1113/jphysiol.2008.164087

Mukund, K., & Subramaniam, S. (2020). Skeletal muscle: A review of molecular structure and function, in health and disease. Wiley Interdiscip Rev Syst Biol Med, 12(1), e1462. 10.1002/wsbm.1462

Roberts, M. D., Young, K. C., Fox, C. D., Vann, C. G., Roberson, P. A., Osburn, S. C., Moore, J. H., Mumford, P. W., Romero, M. A., Beck, D. T., Haun, C. T., Badisa, V. L. D., M. Mwashote, B., Ibeanusi, V., & Kavazis, A. N. (2020). An optimized procedure for isolation of rodent and human skeletal muscle sarcoplasmic and myofibrillar proteins. Journal of Biological Methods, 7(1), e127. 10.14440/jbm.2020.307

Rommel, C., Bodine, S. C., Clarke, B. A., Rossman, R., Nunez, L., Stitt, T. N., Yancopoulos, G. D., & Glass, D. J. (2001). Mediation of IGF-1-induced skeletal myotube hypertrophy by PI(3)K/Akt/mTOR and PI(3)K/Akt/GSK3 pathways. Nature Cell Biology, 3(11). 10.1038/ncb1101-1009

Schmidt, E. K., Clavarino, G., Ceppi, M., & Pierre, P. (2009). SUnSET, a nonradioactive method to monitor protein synthesis. Nature Methods, 6(4), 275–277. 10.1038/nmeth.1314

Tiper, Y., Xie, Z., Hofemeier, A., Lad, H., Luber, M., Krawetz, R., Betz, T., Zimmermann, W. H., Morton, A. B., Segal, S. S., & Gilbert, P. M. (2025). Optimizing electrical field stimulation parameters reveals the maximum contractile function of human skeletal muscle microtissues. American Journal of Physiology - Cell Physiology, 328(4), C1160–C1176. 10.1152/ajpcell.00308.2024

Vandenburgh, H., Shansky, J., Benesch-Lee, F., Barbata, V., Reid, J., Thorrez, L., Valentini, R., & Crawford, G. (2008). Drug-screening platform based on the contractility of tissue-engineered muscle. Muscle and Nerve, 37(4), 438–447. 10.1002/mus.20931

Wilkinson, D. J., Brook, M. S., Smith, K., & Atherton, P. J. (2017). Stable isotope tracers and exercise physiology: past, present and future. Journal of Physiology, 595(9), 2873–2882. 10.1113/JP272277

Wolfe, R. R. (2006). The underappreciated role of muscle in health and disease. The American Journal of Clinical Nutrition, 84(3), 475–482. 10.1093/ajcn/84.3.475

Yeung, S. S. Y., Reijnierse, E. M., Pham, V. K., Trappenburg, M. C., Lim, W. K., Meskers, C. G. M., & Maier, A. B. (2019). Sarcopenia and its association with falls and fractures in older adults: A systematic review and meta-analysis. Journal of Cachexia, Sarcopenia and Muscle, 10(3), 485–500. 10.1002/jcsm.12411552

